# Computer vision to automatically assess infant neuromotor risk

**DOI:** 10.1101/756262

**Authors:** Claire Chambers, Nidhi Seethapathi, Rachit Saluja, Helen Loeb, Samuel Pierce, Daniel Bogen, Laura Prosser, Michelle J. Johnson, Konrad P. Kording

## Abstract

An infant’s risk of developing neuromotor impairment is primarily assessed through visual examination by specialized clinicians. Therefore, many infants at risk for impairment go undetected, particularly in under-resourced environments. There is thus a need to develop automated, clinical assessments based on quantitative measures from widely-available sources, such as video cameras. Here, we automatically extract body poses and movement kinematics from the videos of at-risk infants (N=19). For each infant, we calculate how much they deviate from a group of healthy infants (N=85 online videos) using Naïve Gaussian Bayesian Surprise. After pre-registering our Bayesian Surprise calculations, we find that infants that are at higher risk for impairments deviate considerably from the healthy group. Our simple method, provided as an open source toolkit, thus shows promise as the basis for an automated and low-cost assessment of risk based on video recordings.

## 1 Introduction

Developmental disorders, including those caused by neuromotor disease, are the most common source of childhood disability, affecting 5-10% of children and 3.7 to 7.4 million American children (Rydz, Shevell, Majnemer, & Oskoui, 2005) and are often the cause of lifelong disability. Early intervention may improve outcomes in neuromotor disease, but requires accurate early identification of infants at risk for physical disability (Herskind, Greisen, & Nielsen, 2015; Novak et al., 2017). For effective widespread early identification, easily-deployed automated risk assessment tools are needed to quantify infant movement in the first few months of life during the development of motor control.

There are many factors that describe a ‘good’ diagnostic test. A test’s predictive ability is key. Beyond that, it is desirable that tests are based on quantitative measurements and a series of well-defined steps to reach its conclusion, i.e., an algorithm. This motivates the development of quantitative tests to supplement clinical judgment. Second, it is important to evaluate the availability of a test, or how easy it is to deploy. Clinical assessments often involve expert judgment and expensive equipment, which are only available in highly-resourced environments. This makes assessment inaccessible for families of limited means and in low-resource countries, where the burden of disability is higher (World Health Organization, 2011). Eighty percent of the global prevalence of neuromotor impairment is in low-resource countries, due to larger populations and potentially higher incidence rates (Gladstone, 2010; World Health Organization, 2011). Even in highly-resourced environments, a test that is easy to deploy can enable continuous monitoring of the probability of developing a disorder. Therefore, it is important to assess diagnostic tests not only by their accuracy, but also by how quantitative and cost effective they may be.

Many clinical tests have been developed to assess neuromotor risk. The General Movements Assessment and the Hammersmith Infant Neurological Examination have high sensitivity and specificity. These tests are effective at detecting disorder early, at less than four months corrected age (Novak et al., 2017). The Test of Infant Motor Performance has been shown to detect the changes in the movement patterns pre- and post-treatment for infants later diagnosed with cerebral palsy (Spittle, Doyle, & Boyd, 2008). Despite their high accuracy, these clinical tests have shortcomings. First, they require expert administration and specialized training of licensed clinicians (Bosanquet, Copeland, Ware, & Boyd, 2013; Noble & Boyd, 2012). Therefore, these tests are expensive and only available in well-resourced environments. An automated assessment that objectively quantifies movement characteristics would improve access and reduce costs considerably.

The past two decades have seen the development of sensor-based measurements of infant movement that quantify movement features. Wearable sensors and 3-D motion capture have been used to measure infant movement in laboratory settings (Berg, 2008; Heinze, Hesels, Breitbach-Faller, Schmitz-Rode, & Disselhorst-Klug, 2010; Kanemaru et al., 2014, 2013; Karch et al., 2012; Marcroft, Khan, Embleton, Trenell, & Plötz, 2015; Meinecke et al., 2006; Philippi et al., 2014). Sensor-based measurements are objective and quantitative, and algorithms can generate risk assessments. However, these measurements have been restricted to laboratory and clinical settings, thus hindering the ability to evaluate infants in their natural environments. Moreover, sensor-based methods can be costly and time-intensive to develop and implement, limiting access in resource-poor environments. Thus, sensor-based approaches to diagnostics can provide objective assessment, but have limitations in access and applicability to all infant environments.

In the last decade, scientists have developed systems for video-based assessments using optical flow which hold promise to be widely accessible as they can be implemented on mobile devices (Adde, Helbostad, Jensenius, Langaas, & Støen, 2013; Stahl et al., 2012; Støen et al., 2017). Such methods typically use frame differencing of a video to estimate movements by tracking the centroid of motion. This technique can be extended for the measurement of movement of each limb (Stahl et al., 2012). Using this approach, the amount of movement and the frequency of movement have corresponded to clinical evaluations (Adde et al., 2013, 2010). However, frame difference metrics from optic flow rely on centroid estimates that only measure gross movements as opposed to measuring the kinematic variables for individual joint or limb segment movements. The extraction of limb movement from optic flow also requires careful parameter selection and manual adjustment of the tracking algorithm. As an alternative, marker-less tracking methods have been developed for the tracking of infant movement (Hesse et al., 2018; Olsen, Herskind, Nielsen, & Paulsen, 2014). However, these methods require depth images, are computationally intensive, and the models are likely to be overfit to the relatively small datasets on which they are trained. In sum, video-based diagnostics have the potential to be widely available, but to be successfully applied, they must depend on easily obtained 2-D videos, extract movement features based on individual body-part movements, and require minimal manual tuning.

Here, we provide a first step toward making video-based neuromotor assessment widely available (Figure 1). We produce a ‘normative’ reference database of infant movements using 85 videos found online. We calculate the movements of the body parts using an existing pose-estimation algorithm, OpenPose (Cao, Simon, Wei, & Sheikh, 2017), which we augmented using domain adaptation with our own labelled dataset of infant pose consisting of 9039 infant images. Using this normative database we can calculate how much each infant deviates from the typical movements of healthy infants using a single score, the Naïve Gaussian Bayesian Surprise (Lonini et al., 2016). When we then tested this system on a clinical population (N=19) where the level of neuromotor risk was assessed by a clinician (low, moderate, and high), we found that Bayesian Surprise varied across participant groups. Thus, we have developed an open-source framework that calculates the Bayesian Surprise of movement features given a reference database of ‘normative’ infants.

**Figure 1.**
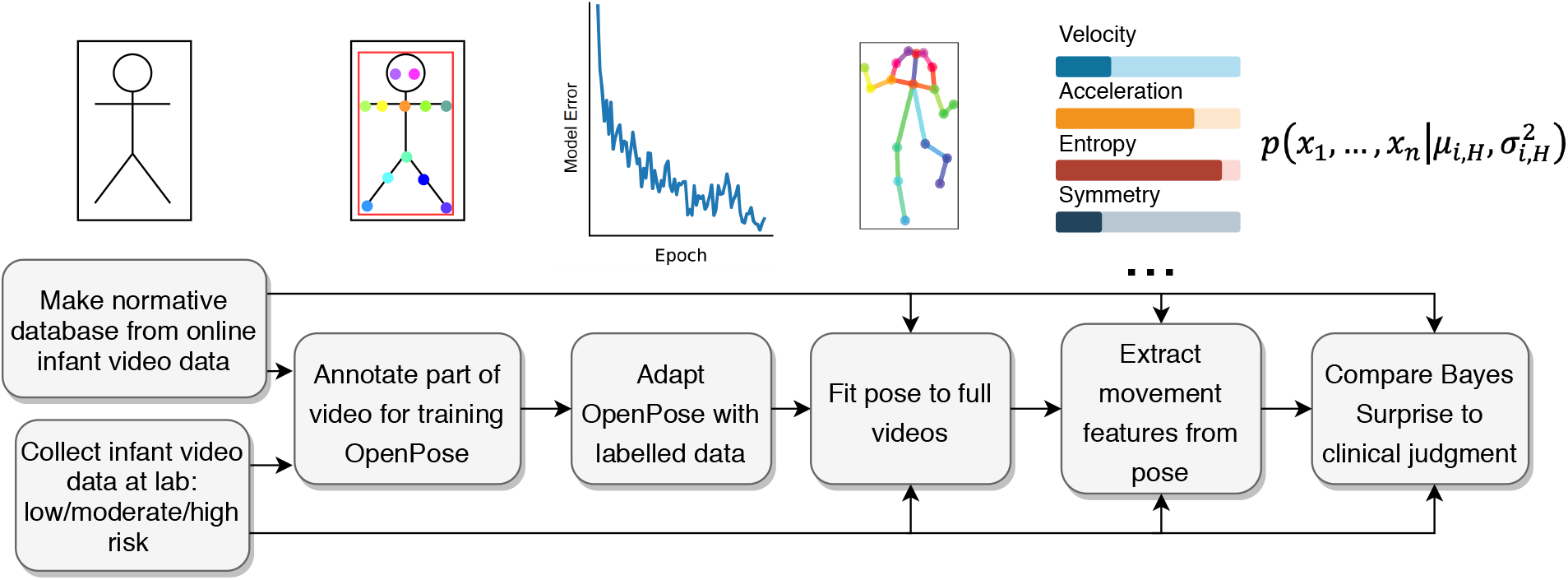
Flowchart of the pipeline for computer vision-based neuromotor risk assessment. We made a normative database infant movement using videos found online (85 infants) and recorded infants at risk of neuromotor disease in a clinical setting (19 infants). Using video frames labelled with body-part landmarks from a subset of our video dataset, we adapted a pose estimator (OpenPose) to extract the pose of infants which we improved using domain adaptation. Using the adapted system, we then extracted pose from all videos. Next, from the pose data, we quantified kinematic features for each infant. Finally, our neuromotor risk prediction used Naïve Gaussian Bayesian Surprise that estimated the probability that each infant belonged to the reference population.

## 2 Method

### 2.1 YouTube Data

In order to develop a framework that predicts the neuromotor risk level of infant movement, it is helpful to have an estimate of what is expected or normal. By creating such a ‘normative’ database, we can then estimate the likelihood that the movements of a new infant are from the normative distribution, thereby assessing the likelihood that the infant is healthy. In previous work we have shown that online video databases are a useful source of human movement data (Chambers, Kong, Wei, & Kording, 2019). In the present study, we built a normative database of infant movements based on video data of infants from YouTube, assuming that found videos represented healthy infant movement. We used search terms such as ‘one-month old’, ‘two-month old’, etc. Search terms that resulted in suitable videos for inclusion in our normative database or for domain adaptation of our pose-estimation system are provided as supplemental data.

In our initial search, we identified 420 video segments featuring infants, where infants were non-occluded, where infants moved independently, and where most of the body was present in the frame throughout the segment. In our analysis, we included 95 of the 420 videos found that were over 5 seconds in duration, and where pose estimates were of sufficient quality for the extraction of basic kinematic variables, as evaluated by visual inspection (mean recording duration per infant = 39s (SD = 33)). Two physical therapists estimated the ages of infants (mean rated age in weeks (SD) = 9.67 (6.26), There was reasonable agreement between raters (inter-rater reliability: *r* = 0.76, *p* < 0.0001, *n* = 85). The age rating averaged across raters was used to classify infants as being more or less than 10 weeks of age (see Data analysis). As supplemental information, we provide URLs and timing information from the YouTube videos in the normative database. When there was more than one video per infant, we included in the dataset the video with the longest duration. This database, containing video recordings of 85 different infants, provided a reasonably large ‘normative’ database of infant movements against which to compare the kinematic features extracted from infant video data collected at the lab.

### 2.2 Clinical Data

Infants tested in person were recruited through The Children’s Hospital of Philadelphia (CHOP), University of Pennsylvania (Penn), and the local community. A subset of the infant data reported here were used in a previous study, which focused on the development of a method to track infant movement using a multiple view stereoscopic 3-D vision system (Shivakumar et al., 2017). Infants were screened to determine eligibility. Inclusion criteria for full-term infants (born at a gestational age of >37 weeks) were the absence of any significant cardiac, orthopedic, or neurological condition. Infants born preterm (gestational age <36 weeks) were recruited from the Newborn/Infant Intensive Care Unit (N/IICU) at CHOP. All infants were between the ages of 3 and 11 months. Infants who could walk were excluded. Parents of eligible participants provided written informed consent. The human subject ethics committee at Penn served as the IRB of record for this study (UPENN IRB # 822487). Data were collected from infants at CHOP, a local child care facility, and at the Penn Rehabilitation Robotics Lab.

Before testing, an experienced pediatric physical therapist evaluated the infants’ level of risk using the Bayley Infant Neurodevelopmental Screener (BINS). This screener includes specific tasks that experts administer to observationally score infants based on the Bayley Scales of Infant Development (Aylward, 1995; Aylward & Verhulst, 2000; Gücüyener et al., 2006). The experts did not have access to the Bayesian Surprise outputs of the algorithm we developed here. We assessed risk of neuromotor dysfunction as classified by the BINS test (low, moderate, high) computed at corrected age for preterm infants and chronological age for full-term infants.

To assess neurodevelopmental risk, the BINS addresses several areas of ability: basic neurological functions/intactness (posture, muscle tone, movement, asymmetries, abnormal indicators); expressive functions (gross motor, fine motor, oral motor/verbal); receptive functions (visual, auditory, verbal); and cognitive processes (object permanence, goal-directedness, problem solving). The BINS consists of six item sets, each containing 11 to 13 items. Each item in the BINS is scored ‘optimal performance’ or ‘non-optimal performance’ using predefined rules. The number of optimal responses for a given item set are summed to give a raw score. For each item set, two previously established raw cut scores identify a given infant’s level of risk for neurological impairment, resulting in three risk groupings: low risk, moderate risk, and high risk.

The Play And Neuro-Development Assessment (PANDA) Gym uses toys with sensors, cameras, and a mat structure which measures the center of pressure of the infant (Figure 2). The video data collected by the gym are pairs of color images extracted from high resolution (1920 × 1080) Go Pro Hero 4 Session videos. Four cameras were mounted on two 3-D printed stereo frames and the videos were captured at 30 frames per second, with one setup placed directly above the baby and the other positioned on the baby’s right side, mounted on the side frame of the gym’s platform. In the current work, we used data from one GoPro camera for each infant. The platform structure is lightweight, made with colored PVC tubing and a sensorized mat (4 × 4 ft), which is developed using a DragonPlate carbon fiber foam core board with four force sensors on each corner. Vinyl and foam padding cover the dragon board to make it comfortable for the infant. Before each infant trial, both systems undergo calibration. A GUI allowed data collection from all sensors and real time tracking of an infant’s center of pressure.

**Figure 2.**
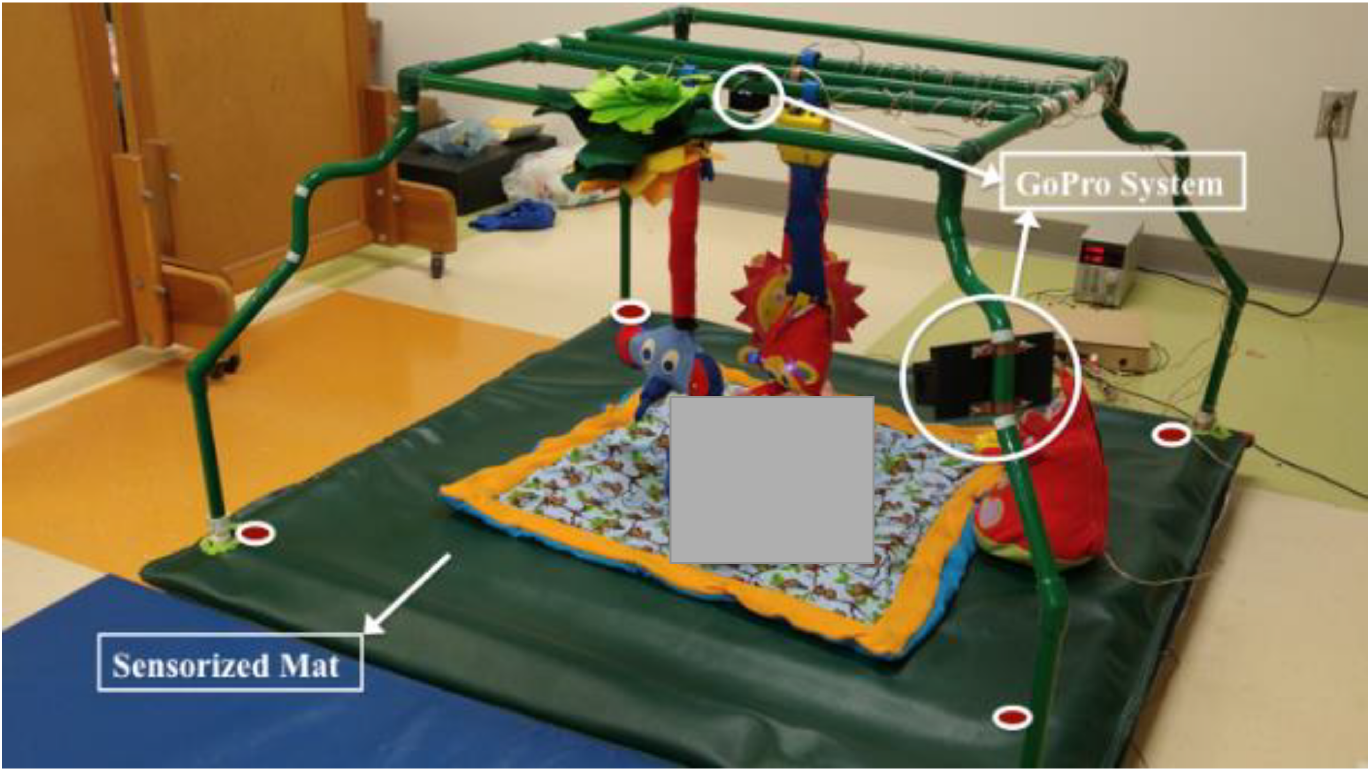
Infant testing at CHOP in the PANDA gym. The infant was placed at the center of a sensorized mat and was recorded using GoPro cameras.

The original purpose of the experiment was to examine the infants’ interactions with sensorized toys, while their movements were recorded using the sensorized mat and GoPro cameras. Sensorized toys contained inertial measurement units (MPU-9150, InvenSense, San Jose, CA) to measure infants’ toy interactions. The experiment included three toy conditions where different toys were hung above the infant (elephant, orangutan, lion, Figure 2) and a fourth baseline condition without the hanging toys (No-toy condition). Since the toys occluded the view of the infant, this caused the pose estimation to fail. Therefore, here, we restricted our analysis to the No-toy condition, where the infant was non-occluded and moved spontaneously. The PANDA gym experiment provided video recordings that allowed us to examine infants’ spontaneous movements.

In each trial, the infant was laid in a supine position inside the PANDA gym and video data were collected using the GoPro cameras. Preterm infants were 2 females and 8 males (mean corrected age (SD) = 14.39 weeks (6.92), mean chronological age (SD) = 24.84 weeks (4.77)). Full-term infants were 17 females and 4 males (mean chronological age (SD) = 26.28 weeks (9.88)). The data of four preterm infants and three full-term infants were excluded due to missing BINS score data. The data of two full-term infants were excluded because infants were sitting during the video, which prevented successful pose estimation. The data of a further three full term infants were excluded due to missing video data.

This left us with a final sample containing 19 infants (Table 1). Preterm infants were 1 female and 5 males (mean corrected age (SD) = 17.64 weeks (4.82), mean chronological age (SD) = 26.74 weeks (4.54)). Full-term infants were 10 females and 3 males (mean chronological age (SD) = 24.65 weeks (9.44)) with 5 infants at low-risk, 9 infants at moderate risk, and 5 infants at high risk, as evaluated by the BINS score. There was a mean (SD) total recording duration of 287.57 s (161.02) per infant.

**Table 1.** Information on infants recorded in PANDA gym included in our analyses. Table includes Infant Id, sex, chronological age, BINS score for chronological age, chronological risk category, corrected age, corrected BINS score, and corrected risk.

| *Infant* | *Sex*<br>*(M,F)* | *Chron.*<br>*age*<br>*(Months)* | *Chron.*<br>*BINS*<br>*score* | *Risk*<br>*(Chron.)* | *Corr. age*<br>*(Months)* | *Corr.*<br>*BINS*<br>*score* | *Risk*<br>*(Corrected)* |
| --- | --- | --- | --- | --- | --- | --- | --- |
| 6 | F | 4.00 | 10 | Moderate |  |  |  |
| 7 | F | 6.75 | 9 | Moderate |  |  |  |
| 8 | F | 4.57 | 10 | Moderate |  |  |  |
| 9 | F | 5.32 | 10 | Moderate |  |  |  |
| 11 | M | 4.00 | 10 | Low |  |  |  |
| 15 | M | 5.78 | 5 | High | 2.50 | 8 | Moderate |
| 16 | M | 8.14 | 0 | High | 5.50 | 0 | High |
| 17 | M | 5.28 | 8 | High | 3.75 | 10 | Low |
| 18 | M | 4.14 | 11 | Low |  |  |  |
| 19 | F | 6.50 | 7 | High |  |  |  |
| 20 | M | 4.50 | 8 | High |  |  |  |
| 23 | F | 11.50 | 7 | Moderate |  |  |  |
| 24 | F | 4.28 | 8 | Moderate |  |  |  |
| 25 | F | 7.75 | 12 | Low |  |  |  |
| 26 | F | 9.50 | 11 | Moderate |  |  |  |
| 27 | F | 7.25 | 11 | Low |  |  |  |
| 30 | M | 7.90 | 3 | High | 5.21 | 8 | Moderate |
| 31 | F | 6.50 | 3 | High | 4.00 | 9 | Low |
| 34 | M | 6.50 | 1 | High | 5.50 | 1 | Moderate |

## 3 Pose Extraction

In order to extract pose information from videos, we used OpenPose, a recently developed pose-estimation algorithm (Cao et al., 2017). OpenPose consists of convolutional neural networks that have been trained using labelled image data to identify 2-D joint and limb positions from images. OpenPose extracts positions of nose, neck, ears, eyes, shoulders, elbows, wrists, hips, knees, and ankles from 2-D videos. On a system with multiple graphical processing units, OpenPose can run in real time (∼ 30 frames/second). With this processing speed, OpenPose provides a viable method for performing motion capture from 2-D videos.

Due to differences between adults and infants in their appearance and pose, pose tracking using OpenPose was initially limited in performance. We therefore adapted OpenPose for infants. We created a dataset of infant images with labels of joint positions. We required that infants be non-occluded, and that their full body be in the frame. We included the first 10s of 103 video segments collected from YouTube and the first 30s of 17 out of the total 19 videos from the clinical dataset. The URLs and timing information of the YouTube video data used for domain adaptation is provided as supplemental data. Videos were selected in order to have a representative set of images for domain adaptation that describe infant pose, in terms of infant appearance, clothing, pose, video background, and image scale. We labelled the video frames with body landmarks using Vatic, a video annotation tool (Vondrick, Patterson, & Ramanan, 2013). Videos were labelled to match the format of the Common Objects in Context dataset (Lin et al., 2014), with labels for left and right eyes, ears, shoulders, elbows, wrists, hips, knees, ankles, and nose, and a bounding box around all points. We included all frames in the labelled dataset, resulting in a total of 36,030 images with labelled pose (32,417 in the training set, 3,613 in the test set). We initialized domain adaptation with pretrained weights from previous work (Cao et al., 2017) and updated weights by gradient descent for 50 iterations. This resulted in a pose estimator that could extract an infant’s pose from 2-D video.

## 4 Data analysis

Using the pose-estimation system adapted for infants, we extracted pose estimates from video recordings of 85 infants in the YouTube cohort and the 19 infants from the clinical sample. From this pose data, we then extracted kinematic features for each infant, then predicted their neuromotor risk using a Naïve Bayes classifier.

When evaluating the pose-estimation model, we used three performance metrics. We computed the root-mean-squared error (RMSE) between ground-truth data and pose estimates, normalized by bounding-box dimensions, for landmarks that were in both datasets. To obtain the RMSE, we computed the distance between each key point detected by the algorithm and label from the ground-truth data, in x coordinates and y coordinates. To account for differences in scale, we normalized x and y distances by the width and height of the ground-truth bounding box around the infant. Finally, we computed the root-mean square of these individual errors from all labelled images, leading to one error score for the entire test dataset. We also evaluated precision, that is, the proportion of total pose estimates that were present in the ground-truth dataset, and the recall, that is, the proportion of ground-truth labels that were found by the pose estimator. These three metrics allowed us to evaluate the pose-estimation model before and after domain adaptation.

Pose-estimation data contained missing data, outliers in the form of false positive detections and additional noise around body landmarks. To obtain clean signals, we preprocessed time series data from each body landmark (Figure 3). We first removed missing data by applying linear interpolation to the raw time series for each body landmark. We then removed outliers by using a rolling-median filter with a smoothing window of 1 second. In order to obtain smooth signals, we then performed smoothing using a rolling-mean filter with a smoothing window of 1 second. Outlier removal and smoothing provided a cleaner signal for the extraction of kinematic variables.

**Figure 3.**
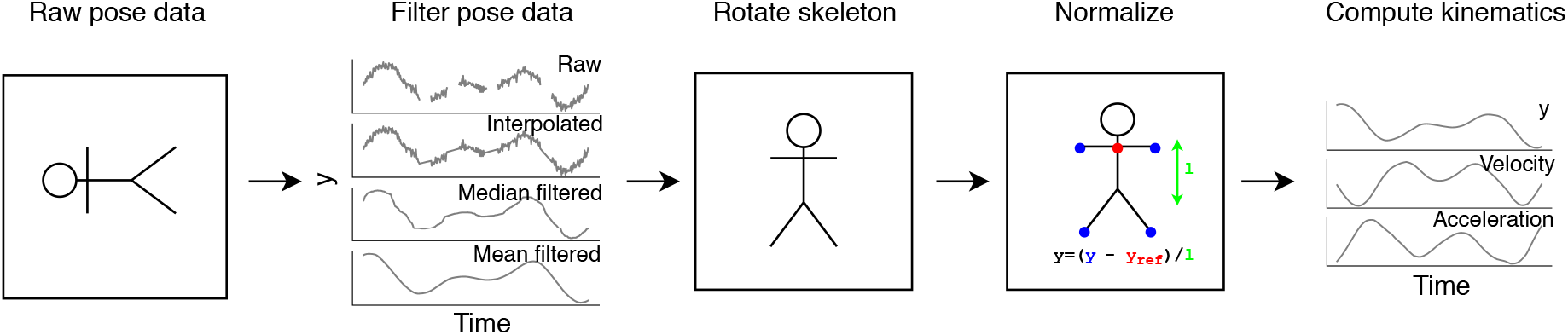
Preprocessing of pose data. We took raw pose data for whole videos as input (frame coordinates of body landmarks). To filter the pose data, we interpolated the raw signal to replace missing data, then applied a rolling-median filter to remove outliers and finally, used a rolling-mean filter. This provided a smooth signal from which to compute derivatives. To ensure that we could compare infants recorded under different conditions (camera angles, video resolution, etc.), we then rotated and normalized body landmark coordinates in each frame. We rotated the upper-body landmarks with respect to the center of the shoulders and we rotated the lower-body landmarks with respect to the center of the hips. Next, we normalized the landmark coordinates within each frame, by subtracting a reference landmark (the neck) and dividing by a reference distance (the trunk length). Finally, based on preprocessed signals, we computed kinematic variables from selected body landmark coordinates and joint angles (position or angle (y), velocity, acceleration).

To compute infant kinematics from YouTube data, we needed to compensate for camera properties. For example, a moving camera with respect to a non-moving infant would produce a movement signal. Also, a short distance between the camera and infant would produce greater movement signals than when further away. To compensate for these differences, we first extracted 2-D joint angles from video frames, which are invariant to the size of the infant in the frame, and the infant’s orientation in the frame (i.e. landscape, portrait). Second, we also computed landmark positions, rotated so that all infants from all frames were aligned (Figure 3). We rotated the upper-body landmarks relative to the angle of the left and right shoulders with respect to the midpoint between the shoulders. We rotated the lower-body landmarks relative to the angle of the left and right hip joints with respect to the midpoint between the hip joints. We then normalized landmark coordinates within each frame (Figure 3). We computed distance from a reference landmark (the neck) and by dividing by a reference distance on the body (trunk length). Measuring movement in body-centered coordinates allowed us to compensate for the effects of camera movement in YouTube videos.

Our predictions of infant risk depended on kinematic variables. From each time series of body-landmark positions, we computed the velocity and acceleration at each time interval. Time-series data extracted from pose estimates provided data for computing kinematic features.

We selected a simple set of features based on movement of the arms and legs computed from knee angles, elbow angles, ankle positions, and wrist positions. Our normalization of pose estimates prevented us from examining movement of the shoulders and hips, so we did not include these as kinematic variables. In future this could be solved by using a fixed camera for all recordings, which would allow us to attribute all recorded movement to the infant and remove the need for normalization. Where possible we computed the median and interquartile range (IQR), to avoid the effects of outliers. We chose features that represented postural information (absolute position and angle), variability of posture (variability of position and angle), velocity of movement (median absolute velocity), variability of movement (variability of velocity and acceleration), complexity (positional and angular entropy), and left-right symmetry of movement (left-right cross correlation of position and angle). As features of each extremity (left and right wrists and ankles), we included the median position in x and y coordinates (units of trunk length, l), IQR of position in x and y coordinates (l), median absolute velocity in x and y coordinates (l/second), IQR of velocity in x and y coordinates (l/second), IQR of acceleration in x and y coordinates (l/second^2^), left-right cross-correlation of position, and positional entropy. As features of each joint angle (left and right elbows, wrists, knees and ankles), we included the angular mean (degrees), angular standard deviation (degrees), median angular velocity (degrees/second), IQR angular velocity (degrees/second), IQR angular acceleration (degrees/second^2^), angular cross-correlation, and angular entropy. To reduce the number of features, we averaged feature values across the left and right sides of the body, where applicable. Having established a basic set of features, we pre-registered our feature set and the algorithm to calculate Bayesian surprise (https://osf.io/hv7tm/). Our preregistered set of kinematic features provided basic descriptors of infant movement on which to make predictions.

Existing assessments rely on clinicians visually identifying at-risk infants whose movements differ from the normal population. We aimed to replicate this form of assessment using quantification of multiple kinematic features from videos and a normative database which tells us what healthy infant movement looks like. In this approach, infants who deviate from the healthy reference population are identified as being at-risk. We estimated the probability of each infant belonging to the healthy reference population, represented by the YouTube cohort in this case, who are presumed to be healthy. We adopted the Naïve Bayes approach, previously used for clinical assessments of walking performance (Lonini et al., 2016). Under the assumption of normally-distributed features and feature independence, the joint probability of features over the reference population is:

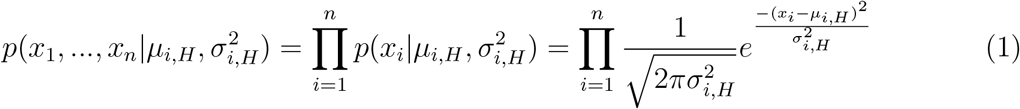

where *x*_*i*_ indicates the *i*-th feature value for a subject, and *µ*_*i,H*_ and 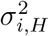 are, respectively, the mean and variance of that feature across the reference subjects.

The negative natural logarithm gives the Naïve Gaussian Bayesian Surprise:

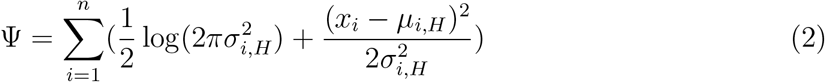

This metric can be interpreted as being related to the log probability of a subject being part of the reference population. The Naïve Bayesian Surprise was normalized with respect to the reference population. Infants were compared within age brackets computed based on corrected age (less than 10 weeks, 10 weeks or older). We chose 10 weeks as a threshold so that there was sufficient data within each age bracket to serve as a normative group for comparison (47 infants less than 10 weeks, 38 infants older than 10 weeks). This provided a standard score: subjects who were more than two standard deviations from the mean would be classed as at risk. Our approach to risk assessment classified infants based on how different their movement was to that of the reference population.

To further explore how combinations of movement features relate to clinically-assessed risk, we applied matrix decomposition to our set of 38 movement features from 104 infants. Matrix decomposition provides a way of describing infants’ movement in terms of a lower number of features which are aggregates of the original set of 38 movement features. This method also allows us to summarize relationships between movement features. Singular value decomposition (SVD) is a matrix-decomposition method that finds ‘latent variables’ which describe important sources of variance in the data. Each latent variable is a linear combination of the original set of movement features, with a singular value that tells us how much of the variance it explains. For example, one latent variable could have a high weight for velocity and acceleration, and a low weight for symmetry and entropy; a second latent variable could have a high weight for symmetry and entropy, and a low weight for the remaining features, and so on. These groupings of features into one latent variable tell us that they vary together. Each infant has one score for each of the latent variables. For example, an infant with high velocity and acceleration, and moderate symmetry and entropy would have a high score for the first latent variable and a moderate score for the second latent variable. Thus, matrix decomposition allows description of infants’ movement in terms of a lower number of features.

More formally, applied to our dataset of *M* infants by *N* movement features (*A*_*M*×*N*_), SVD approximates this matrix by the product of three matrices: *A* = *U* Σ*V*^*T*^. The columns of the left matrix, *U*, span the column space of matrix *A* that characterizes the infants, later described as singular vectors. The columns of the right matrix, *V*, span the row space of matrix *A* that characterizes the movement features, later described as singular vector weights. Σ contains singular values that describe how important each latent variable is. We first normalized columns of matrix *A*, by subtracting the mean and dividing by the standard deviation. We then applied SVD. We examined the singular vectors of *U* as a function of risk and we examined how the original movement features are weighted in latent variables by examining singular vectors of *V*. SVD allows examination of infants and movement features in terms of a set of latent variables that describe the sources of variance in the dataset.

## 5 Results

We developed a system to assess infants’ neuromotor risk based on kinematic variables extracted from 2-D videos. In order to estimate the pose of infants from videos, we first adapted an existing pose-extraction algorithm for infants using data on infant pose. We compared kinematic data extracted from infants at risk of neuromotor disorders (low, moderate, high risk) with a normative database from videos of 85 infants. Based on pose estimates, we quantified basic kinematic features of movement. We combined these features into a single estimate of risk by computing Bayesian surprise for each infant in a cohort of 19 infants seen in a clinical setting.

We required a system that could reliably extract the pose of infants from videos. We therefore adapted an existing deep-learning based pose-estimation algorithm, OpenPose (Cao et al., 2017), for infants. We performed domain adaptation for infant pose extraction by updating the network’s weights with infant images and body landmark labels as input. Domain adaptation led to improved pose estimation in infants. We observe a lower distance between ground-truth labels and pose estimates after domain adaptation, as shown by the RMSE in bounding-box units. RMSE of 0.05 before domain adaptation decreases to 0.02 after domain adaptation (Figure 4 A,B). We also observe an increase in precision from 0.89 before domain adaptation to 0.92 after domain adaptation (Figure 4C,D) and an increase in recall from 0.76 before domain adaptation to 0.94 after domain adaptation (Figure 4E,F). Therefore, through domain adaptation with labelled images of infants, we improved the performance of the pose-estimation model. This allowed us to extract skeletal information on each infant and track the positions of body landmarks (Figure 4 G,H). The adapted pose estimator allowed us to extract movement trajectories from videos of infants moving.

**Figure 4.**
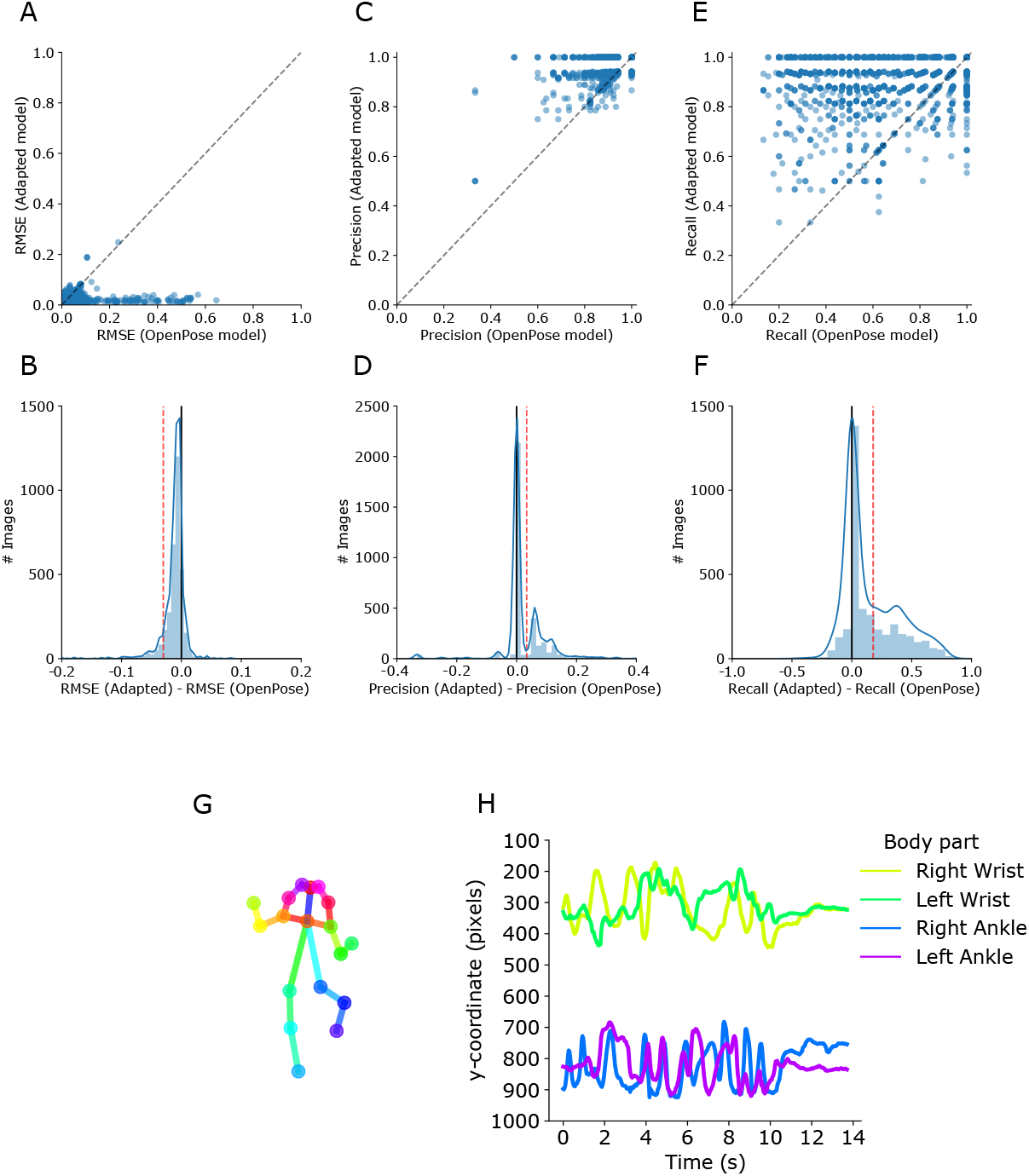
Pose-estimation model performance and example OpenPose outputs. (A) Scatterplot of the RMSE of the adapted pose-estimation model as a function of the RMSE of the OpenPose model before domain adaptation. Points show the RMSE for individual images in bounding-box units. In A, C and E, the dotted line shows the diagonal, where performance before and after domain adaptation are equal. (B) Distribution of the difference in model error before and after domain adaptation. For each model, a single RMSE score was computed from errors between individual key points and labels averaged across the whole test dataset. RMSE (Adapted) – RMSE (OpenPose) is shown by the red dotted line. Solid black line shows a difference of 0, in B, D and F. The negative RMSE difference demonstrates improvement after domain adaptation. (C) Scatterplot of the precision of the adapted pose-estimation model as a function of the precision before domain adaptation. Points show the precision for individual images. (D) Distribution of the difference in precision before and after domain adaptation. Precision (Adapted) – Precision (OpenPose) is shown by the red dotted line. (E) Scatterplot of the recall of the adapted pose-estimation model as a function of the recall before domain adaptation. Points show the recall for individual images. (F) Distribution of the difference in recall before and after domain adaptation. Recall (Adapted) – Recall (OpenPose) is shown by the red dotted line. (G) Example of OpenPose outputs extracted from a YouTube video of an infant using our adapted pose-estimation system. (H) Image y-coordinates of the extremities for the same infant as in (G).

We compared the movement of each infant from the clinical cohort with the movement extracted from the reference-group sample based on 38 features that described posture, velocity, acceleration, left-right symmetry, and complexity (see methods, all pre-registered). Inspection of kinematic features from different risk populations reveals some subtle deviations from the reference sample based on neuromotor risk (Figure 5). While each of these features may not be strongly indicative of an infant’s neuromotor risk when considered individually, an estimate which pools across features is likely to provide more robust predictions of risk.

Our approach to assessing neuromotor risk allowed us to combine many features into one estimate. We computed the Bayesian Surprise for each individual infant. When features are combined into one estimate of risk, the normalized Bayesian Surprise increases with clinician-assessed risk; Low Risk: mean z=−1.62, SD=1.18; Moderate Risk: mean z=−1.68, SD=1.34; and High Risk: z=−2.94, SD=1.43 (Figure 6). The proportion of infants classed as at risk varies with participant group (Reference: P(Risk) = 0.07, Low Risk: P(risk) = 0.40, Moderate Risk: P(Risk) = 0.33, High Risk: P(Risk) = 0.8). A Kruskal-Wallis test showed a significant association between participant group and the Bayesian Surprise score (*χ*^2^(3) = 29.92, p<0.0001). We found significant differences between the reference population and the clinical risk groups using the Mann Whitney U test (Reference, Low Risk: U = 49, p<0.05; Reference, Moderate Risk: U = 108, p<0.005; Reference, High Risk: U = 9, p<0.005, Bonferroni-corrected p-values). Differences among clinical risk groups were non-significant (Low Risk, Moderate Risk: U = 20, p=0.99; Moderate Risk, High Risk: U = 6, p=0.63; Low Risk, High Risk: U = 12, p=0.54, Bonferroni-corrected p-values). These proof-of-concept results show differences between at-risk infants and our reference population based on a set of simple kinematic features. These results suggest that it may be possible to predict clinician’s assessments of infantile neuromotor risk through statistical comparison relative to a healthy reference population.

**Figure 5.**
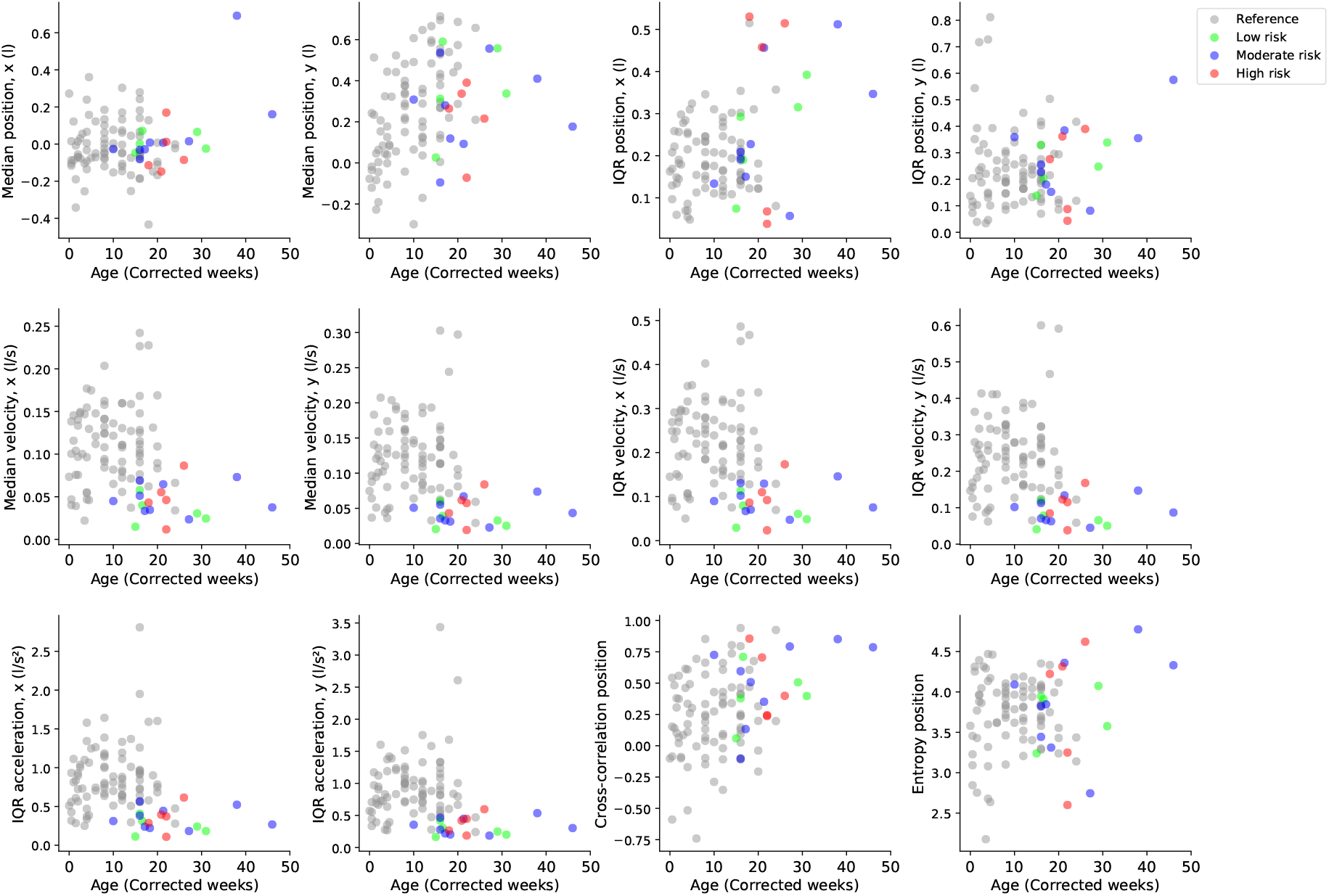
Infant movement features. Kinematic features of the reference sample (gray), low-risk infants (green), moderate-risk infants (blue), high-risk infants (red) as a function of age in corrected weeks. Features are shown for the wrists: median absolute position (l), IQR of position (l), median velocity (l/s), IQR of velocity (l/s), IQR of acceleration (l/s^2^), left-right cross-correlation of position and entropy of position. Visualization of other features are provided as Supporting Information.

**Figure 6.**
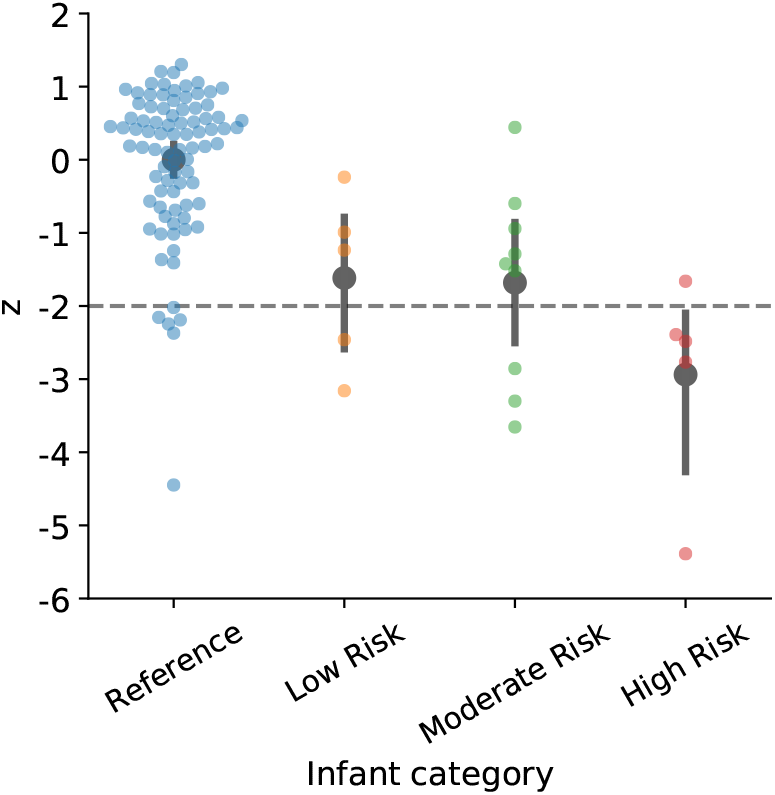
Normalized Bayesian Surprise as a function of subject group. The normalized Bayesian Surprise (z) is shown for the reference infant population, and at-risk infants recorded at the lab evaluated by clinicians using the BINS score (Low Risk, Moderate Risk, and High Risk). More negative scores indicate a smaller probability of belonging to the reference population, or higher risk. Points show individual data for each subject group. Individual data is overlaid with the mean for each group (error bars = 95% confidence intervals (CI)).

We explored how combinations of our set of 38 movement features relates to neuromotor risk. SVD finds linear combinations of movement features, or latent variables, that summarize the movement feature data using matrix factorization. In our data, the three most important latent variables account for 37%, 15%, and 12% of the variance respectively (Figure 7). We did not include other variables, since they accounted for small proportions of variance (<10%). The first latent variable mainly discriminates between the reference group and infants recorded at the lab, with at-risk infants being more extreme along this variable (Figure 7A). Well-represented movement features include median and variability of velocity, and variability of acceleration (Figure 7B). The second latent variable discriminates between reference, low-medium risk infants and high-risk infants, with high-risk infants having more extreme values along this variable (Figure 7C). Variability of position, positional entropy, symmetry (cross-correlation), and postural variables from the lower body (mean knee angle, median ankle position) are well-represented (Figure 7D). Therefore, in our data, high-risk infants are atypical for this combination of features. The third latent variable does not show a clear pattern relative to the infant group (Figure 7E), but may relate to movement of the upper body relative to the lower body, as most positive weights describe lower body movement and most negative weights describe upper body movement (Figure 7F). Thus, our exploratory analysis suggests that at-risk infants may move more slowly, and have less variable velocity and acceleration, and that high-risk infants in particular may have more extreme posture, variability of posture, and symmetry. These results show that at-risk infants in our sample have different movement patterns to the normal population, which can be described in terms of combinations of individual movement features. This analysis allows us to describe important sources of variance in the movement feature data and to explore the importance of combinations of movement features in determining risk assessments.

**Figure 7.**
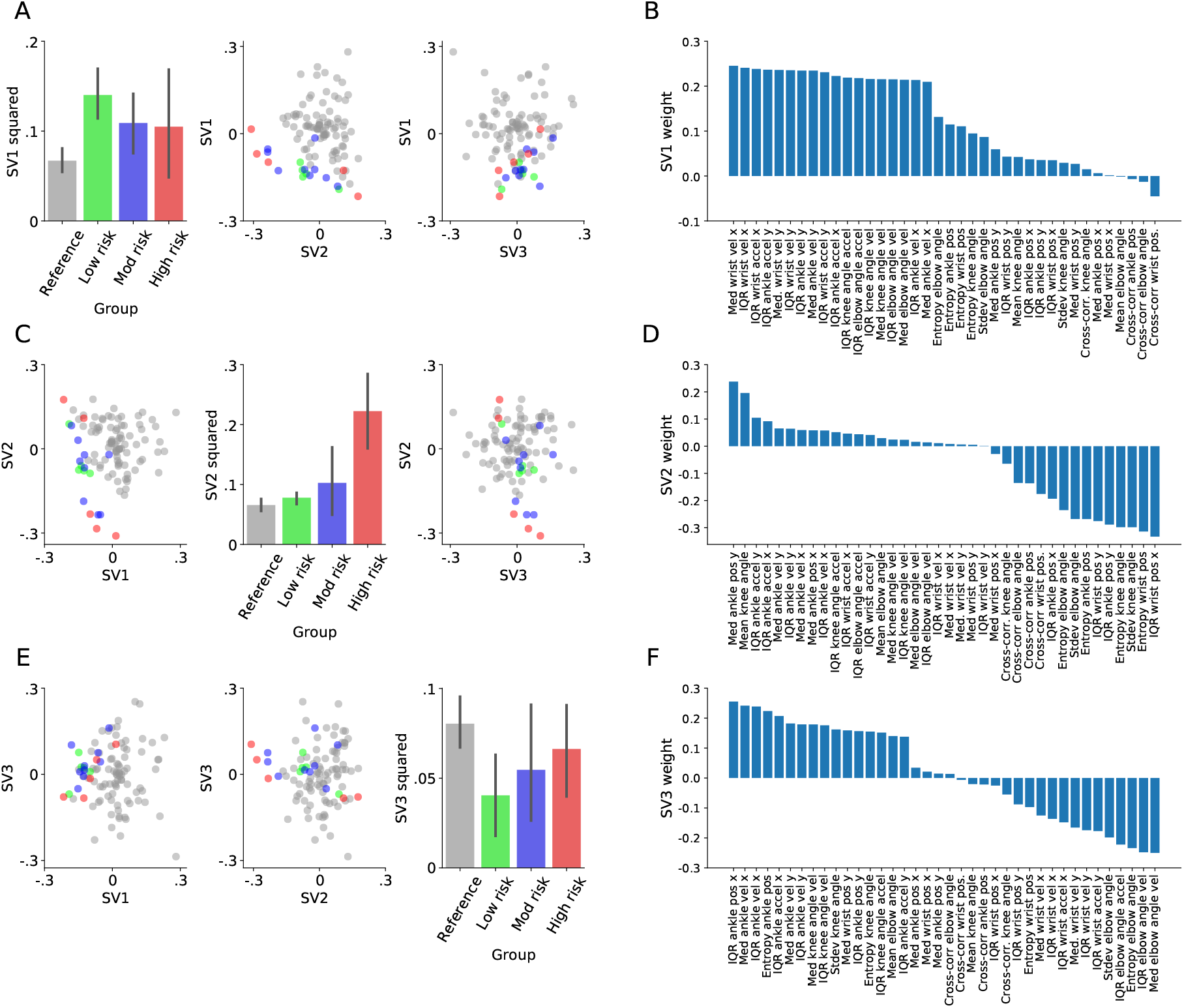
SVD analysis of movement feature data in terms of the three most important latent variables. (A), (C), and (E) show values from singular vectors, which describe the infants in terms of latent variables. (B), (D), and (F) show the weighting of movement features in each latent variable. (A) Left: Mean eccentricity (error bars=95% CI) along the first singular vector (SV1 squared), as a function of participant group. Center, Right: Scatterplots displaying values of the first singular vector as a function of the second and third singular vectors (SV2, SV3) for individual infants. (B) Weighting of movement features in SV1 ranked in descending order. (C) Left, Right: SV2 as a function of the SV1 and SV3 for individual infants, with participant shown by color. Center: Mean SV2 squared (error bars=95% CI) as a function of neuromotor risk. (D) The weighting of movement features in SV2 ranked in descending order. (E) Left, Center: SV3 values as a function of the SV1 and SV2 for individual infants. Right: Mean SV3 squared (error bars=95% CI) as a function of neuromotor risk. (F) Weighting of movement features in SV3 ranked in descending order.

## 6 Discussion

We have developed a framework to identify infants at risk of neuromotor disorder based on video data. Our approach was to compare infants assessed in the laboratory at different levels of neuromotor risk with a normative dataset of infant movement extracted from online videos. We successfully adapted an existing pose-estimation system to perform motion capture on 2-D videos. Using pose estimation, we extracted features from videos that reflected posture, kinematic variables (velocity, acceleration and their variability), complexity (entropy), and symmetry (left-right cross correlation). We found that the Bayesian Surprise measure of risk which pooled across features varies across participant groups. We examined the main latent variables that describe the data, with the finding that velocity and acceleration distinguish infants from the reference population with at-risk infants, and that high-risk infants differ from normal in their posture, postural variability, and symmetry. Combinations of movement features are predictive of neuromotor risk.

Previous work has addressed kinematic markers of disordered infantile movement, with the finding that features of velocity and acceleration including skewness of velocity, maximum acceleration, and minimum speed predict infantile movement disorders (Heinze et al., 2010; Meinecke et al., 2006). Other work has shown that infants with disordered movement show greater stereotypy (Karch et al., 2012; Philippi et al., 2014). Our SVD analysis showed that at-risk infants may have lower velocity (median and IQR) and variability of acceleration relative to a reference sample of healthy infants, and that high-risk infants in particular show different patterns of posture, postural variability, symmetry of movement, and complexity. Our findings on the features that predict risk assessments show some similarity with previous observations with roles for velocity and acceleration, as well as complexity and symmetry. In order to confirm our findings, it will be important to examine the kinematic markers of disordered movement using large clinical datasets.

Previous work on infant movement used measurements capable of extracting fine movement information, like accelerometers, motion capture using depth cameras, and optic flow from videos (Adde et al., 2010; Heinze et al., 2010; Meinecke et al., 2006). This allowed examination of the jerkiness of movement, previously found to be predictive of movement disorder (Heinze et al., 2010; Meinecke et al., 2006). Pose estimation applied to low-quality videos currently does not afford the high precision needed to compute higher derivatives of movement trajectories. Here, because of the presence of outliers and noise in our pose-estimate data, we applied smoothing to the data which removed its fine detail. In future work, improvements in pose estimation and high-quality video datasets will make it possible to extract fine movement detail from pose estimates and will allow examination of more subtle movement features.

One major concern about our work here is the small number of subjects in the clinical group. Due to a range of constraints we were only able to analyze data from 19 subjects. We do find a significant difference of those subjects to the normative population. However, we find a big difference even when comparing the normative population to the low-risk infants in the clinical cohort. Our SVD-based analysis (Figure 7) provides some insights into why we are seeing these changes. Features including velocity, acceleration and their variability discriminate between at-risk and reference infants. Future work will need to correct for sources of variability which do not directly correspond to neuromotor risk. While our analysis was exactly as we had planned and preregistered, we cannot avoid the confound that other differences between the two groups gave rise to the difference we observed. As such, our study only provides a low level of certainty and should be followed up with a considerably larger clinical population.

In this work, we provided methods to quantify and automate infantile neuromotor risk assessments. This system could benefit from several additions. For example, infants move differently depending on the time of day and their emotional state (Oyerinde et al., 2018). For an automated movement test with high enough accuracy to be used, it will be important to take such variables into account. It is also important to take into account developmental changes in infant movement (Law, Lee, Hülse, & Tomassetti, 2011). Development is variable across infants: an infant can be delayed in their motor development without having a neuromotor disorder (WHO Multicentre Growth Reference Study Group, 2006). Therefore, an algorithm that jointly infers an infant’s developmental age and neuromotor risk promises to perform better than one that infers risk alone. The incorporation of several variables will be needed for high-accuracy predictions of neuromotor risk.

Our approach to infantile neuromotor risk assessments has made novel contributions. For our normative database, we collected movement data from close to 100 infants using marker-less tracking applied to videos. Use of a normative database for comparison increases the robustness of risk assessments. Secondly, we used unsupervised methods to assess risk and to examine important movement features. The Bayesian Surprise metric compared movement of each at-risk infant to the normative database, weighting each feature by its uncertainty. Our SVD analysis describes the main sources of variance in the movement feature dataset. Therefore, our analysis is unlikely to be overfit to the data.

Not only does marker-less tracking allow access to larger infant kinematic datasets, but also allows assessments to be based on movement under conditions outside clinical settings. This is advantageous for two reasons. First, video-based diagnostics promise improved access to evaluations. A parent need only provide a simple smartphone video of their infant’s movement to receive an assessment. Second, as with experimental paradigms, behavior observed in the laboratory and clinical setting may differ from the real-life setting (Bronfenbrenner, 1977; Ingram & Wolpert, 2011). Movement variables collected in natural settings would provide a more ecologically-valid assessment, thus providing more accurate predictions.

Marker-less tracking not only has strong potential for applications, but also as a tool for understanding infant pathology. One of the most common movement disorders is cerebral palsy, a lifelong condition due to brain injury in infancy. Although subtypes and biomarkers have been described, it is poorly understood in terms of its causal determinants (Sanger, 2008). Marker-less tracking may serve as a tool to provide a detailed description of movement pathology across a large population, providing better quantitative descriptions of disease subtypes (Katz & Rymer, 1989; Rosenbaum et al., 2007; Sanger et al., 2006). Based on large-scale quantitative measurements of movement in different patient groups and in typical populations, one could model how motor-control processes differ between healthy and disordered infants (Scott & Norman, 2003). Therefore, marker-less tracking also provides a tool for understanding infant pathology.

We have developed a method to identify infantile risk of neuromotor disorder based on pose estimates extracted from 2-D videos. Such a method meets the requirements of objectiveness, as movements are assessed based on quantitative variables; and availability, since diagnoses would no longer require the opinion a trained specialist. Our approach proposed here will improve as larger normative datasets are collected, and as pose-estimation algorithms better suited to movement science are developed (Seethapathi, Wang, Saluja, Blohm, & Kording, 2019), allowing predictions to be made from combinations of more subtle movement features. We expect that, over the next decade, movement-based diagnostics from pose estimates will become a viable alternative to established tests such as the General Movements Assessment (Prechtl, 2001; Wei & Kording, 2018).

## 6.1 Supporting information

Code and data referenced in the manuscript are provided at: https://github.com/cchamber/Infant_movement_assessment/

https://doi.org/10.6084/m9.figshare.8161430

## 6.2 Acknowledgements

We thank members of Professor Michelle Johnson’s laboratory, in particular Wilson Torres, who helped with labelling image data. We also thank Professor Kunlin Wei and lab members, Dr. Gaiqing Kong and Xiaoyue Wang, who helped with both labelling image data and with curation of infant video data from YouTube. We thank Dr. Frances Shofer for helpful input and comments.

## Notes

https://doi.org/10.6084/m9.figshare.8161430

